# In silico analysis of single nucleotide polymorphisms (SNPs) in human C-C chemokine receptor type five (CCR5) gene

**DOI:** 10.1101/2020.11.14.382739

**Authors:** Asma Ali Hassan Ali, Muntaser Ibrahim

## Abstract

**Introduction:** Chemokines are small transmembrane proteins with immune surveillance and immune cell recruitment functions. the expression of CCR5 gene affects virus production and viral load(1). The CCR5 gene contains two introns, three exons, and two promoters, and it is necessary as a co-receptor for the entry of the macrophage-tropic HIV strains. Mutations in the coding region of CCR5 affect the protein structure, which will affect production, chemokine binding, transport, signaling and expression of the CCR5 receptor.

**Methods:** SNPs within CCR5 gene were retrieved from ensemble database. Coding SNPs were analyzed using SNPnexus. Coding non-synonymous SNPs in CCR5 binding domains with Viral gp120 were analyzed using SIFT, PolyPhen and I-mutant tools. Project HOPE then used to modelled the 3D structure of the protein resulting from these SNPs. Non-coding SNPs that affects miRNAs in 3’ rejoin were analyzing using PolymiRTS. SNPs that affect transcription factor binding were analyzed using regulomeDB.

**Results:** (178) non-synonyms missense SNPs were found to have deleterious and damaging effect on the structure and function of the protein. In CCR5 binding domains with Viral gp120: 3 SNPs rs145061115, rs199824195 and rs201797884 were found to affect both structure and function and stability of chemokine protein. The 2 SNPs rs185691679 and rs199722070 has a role in disruption and creation of the target sites in miRNA seeds due to their high conservation score.

**Conclusion:** Mutations in CCR5 gene may explain and represent the molecular basis of the resistance to HIV infection.

## 1. Introduction and literature review

Chemokines are small transmembrane proteins with immune surveillance and immune cell recruitment functions. The effects of these chemokines are mediated by their G-protein-coupled receptors (GPCR’s), which upon binding to the relevant ligand results in the release of the αi and βγ G-protein subunits. Other chemokine receptors work together with CCR5 to stimulate T-cell functions(2). This, in turn, mediates an effector response (1, 3). Depending on the structure and number of cysteine residue Chemokines are classified as C, CC, CXC, and CX3C. The C-C receptors often share a significant degree of homology, with ‘C-C chemokine receptor type five’ (CCR5) and ‘C-C chemokine receptor type two’ (CCR2) sharing 75% homology (4). The chemokine receptor CCR5 is expressed on various cell populations including macrophages, dendritic cells and memory T cells in the immune system, endothelium, epithelium, vascular smooth muscle and fibroblasts, and microglia, neurons, and astrocytes in the central nervous system(5). In 1996, it was discovered that CCR5 is necessary as a coreceptor for entry of the macrophage-tropic HIV strains(6, 7). In the Structure of CCR5 coreceptor, the HIV binding domains are a N-terminal and ECL2 domain, the ECL1 domain is indicated. Viral gp120 has been shown to bind sulfate moieties on the cell surface (8, 9). During initial infection, the virus uses CCR5, whereas the alternative coreceptor ‘C-X-C chemokine receptor type four’ (CXCR4) is used in later HIV infection when the infected individual is progressing to AIDS.

The CCR5 protein consists of 352 amino acids with a molecular weight of 40.6 kDa (10). The protein consists of amino-terminal (N-terminal), three extracellular loops (ECL), three intracellular loops (ICL), cytoplasmic or carboxyl tail (C-terminal tail) and seven transmembrane domains (TMD) made up of hydrophobic residues. These hydrophobic regions are important to chemokine ligand binding, HIV coreceptor activity and functional response of the receptor. The N-terminal is rich in tyrosine and acidic amino acids which facilitating interaction with ligands and HIV(11). The I12T, C20S and A29S variants are all located in the N-terminal. the variants markedly reduce cell surface expression and ligand binding with HIV co-receptors(12). The C20S variant prevents disulfide bond formation between the N-terminal and ECL 3. Considering the importance of this bond in chemokine binding(13). The CCR5 gene was localized in chromosome 3p21(14) and was found within a cluster of genes encoding for other chemokine receptors which included CCR1, CCR2, CCR3, XCR1 and CCBP2(10, 15). The CCR5 gene is composed of three exons, two introns, and two promoters. The two promoters for CCR5, Pu and Pd contain several ATG transcription sites, prior to the start codon of exon 3, leading to the generation of different CCR5 transcripts, which vary in their 5’ UTR regions (16). Mutations in the Coding region of CCR5 affect the protein structure, which will affect production, chemokine binding, transport, signaling and expression of the CCR5 receptor. Mutations in the promoter region will affect the DNA transcription factor (TF) binding or regulatory sites leading to change in mRNA production of CCR5.

The CCR5Δ32 mutation was discovered as a genetic mutation that protect the cells from infection by HIV (7, 14, 17), The deletion involves a frameshift mutation result in mutant allele contains 215 amino acids in comparison to the full-length 352 amino acid wild type CCR5.the affected rejoin was the second extracellular loop(17), The subsequent protein lacked the last three transmembrane domains as well as regions important in G-protein interaction and signal transduction. The Δ32 mutant allele is confined mostly to individuals of European descent, at gene frequencies of approximately 10%, and has a north to south latitude decline in frequency(18).

In this study, we performed computational analysis of all SNPs in the CCR5 gene found in ensemble database in order to identify coding and non-coding SNPs that can possibly modify the structure and function of chemokine receptor.

## 2. Materials and methods

All SNPs and their related protein sequences within the CCR5 gene were retrieved from ensemble (https://www.ensembl.org/index.html)(19) and uniport (https://www.uniprot.org/) (20) database respectively. We used “CCR5” as our search term in the ensemble and used filters to narrow down our search results into three categories: coding region SNPs, non-coding region SNPs and SNPs in the CCR5 binding domain with Viral gp120. In coding region, the deleterious nsSNPs were detected by the web program SIFT and PolyPhen both in SNPnexus (http://www.snp-nexus.org/) (21). SIFT was used to distinguish between tolerant and intolerant coding mutations and is used to predict whether an amino acid substitution in a protein will have a phenotypic effect. PolyPhen is a computational tool for identification of potentially functional nsSNPs. We took the SNPs in binding domains predicted by both databases SIFT and PolyPhen to be deleterious and damaging (3 SNPs; rs145061115, rs199824195 and rs201797884) to Predict the molecular phenotypic effects of these SNPs using Project HOPE (http://www.cmbi.ru.nl/hope/)(22). It is an online webserver used to search protein 3D structures with explanation of such a change in both structure and function of the protein, we entered the three SNPs (rs145061115, rs199824195, and rs201797884) that we obtained from the last step with the primary structure of the CCR5 protein (obtained from uniport) into HOPE. I-mutant 2.0 tools (http://folding.biofold.org/i-mutant/imutant2.0.html) was used to predict the effect of these substitutions on the stability of chemokine receptor protein. For Characterization of SNPs in non-coding region, we used regulome web server (http://www.regulomedb.org/) to predict the functional effect on transcription factors binding which might may affect the level, location, or timing of gene expression. And also, we used PolymiRTS v3.0 (http://compbio.uthsc.edu/miRSNP/) to predict the functional impact of these SNPs in miRNA seed regions and miRNA target sites which might lead to a decrease/increase of the expression of CCR5 protein. The analysis steps of the functional assessment of in-silico processes are summarized in Figure 1.

**Figure 1:**
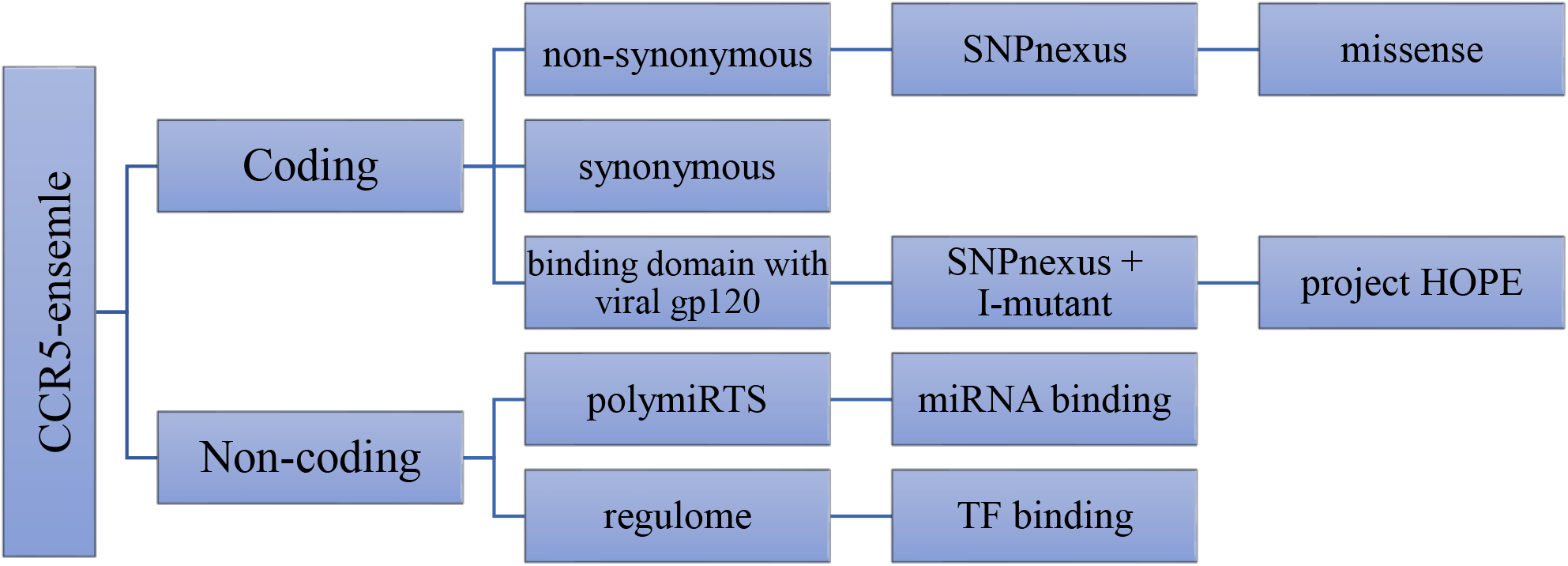
Flowchart of the analysis processes.

## 3. Results

Among two thousand five hundred and ninety (2590) SNPs within CCR5 gene retrieved from ensemble database we selected SNPs in coding and non-coding region, 960 are in coding region including 259 synonyms, 701 non-synonymous, ((40) frameshift, (661) missense).in the prediction of coding nsSNPs, 178 are found to be deleterious and damaging, 37 are nonsense (gain stop codon) SNPs. In a total of 1630 non-coding region SNPs, 507 are intronic, 665 in 5’and 3’ UTR, 489 are in upstream region and the rest four hundred thirty-nine (439) are downstream SNPs.

**Table1:** Three SNPs in the CCR5 binding domain with Viral gp120 (rs145061115, rs199824195, and rs201797884) which correspond to C20S, C178R and (W190R/W190G) respectively are found to affect the structure and function and stability of chemokine receptor protein predicted by SIFT, PolyPhen, and I-mutant tools.

| SNP ID | Location | Alleles | AA change | SIFT _ class | SIFT score | polyphen _class | Poly-phen score | stability | Reliability Index (RI) |
| --- | --- | --- | --- | --- | --- | --- | --- | --- | --- |
| rs145061115 | 3:46372960 | T/A | C20S | deleterious | 0.01 | probably damaging | 0.994 | Decreased | 7 |
| rs199824195 | 3:46373434 | T/C | C178R | deleterious | 0 | probably damaging | 1 | Decreased | 6 |
| rs201797884 | 3:46373470 | T/C/G | W190R | deleterious | 0 | probably damaging | 0.999 | Decreased | 8 |
| rs201797884 | 3:46373470 | T/C/G | W190G | deleterious | 0 | probably damaging | 0.999 | Decreased | 9 |

**Figure2:**
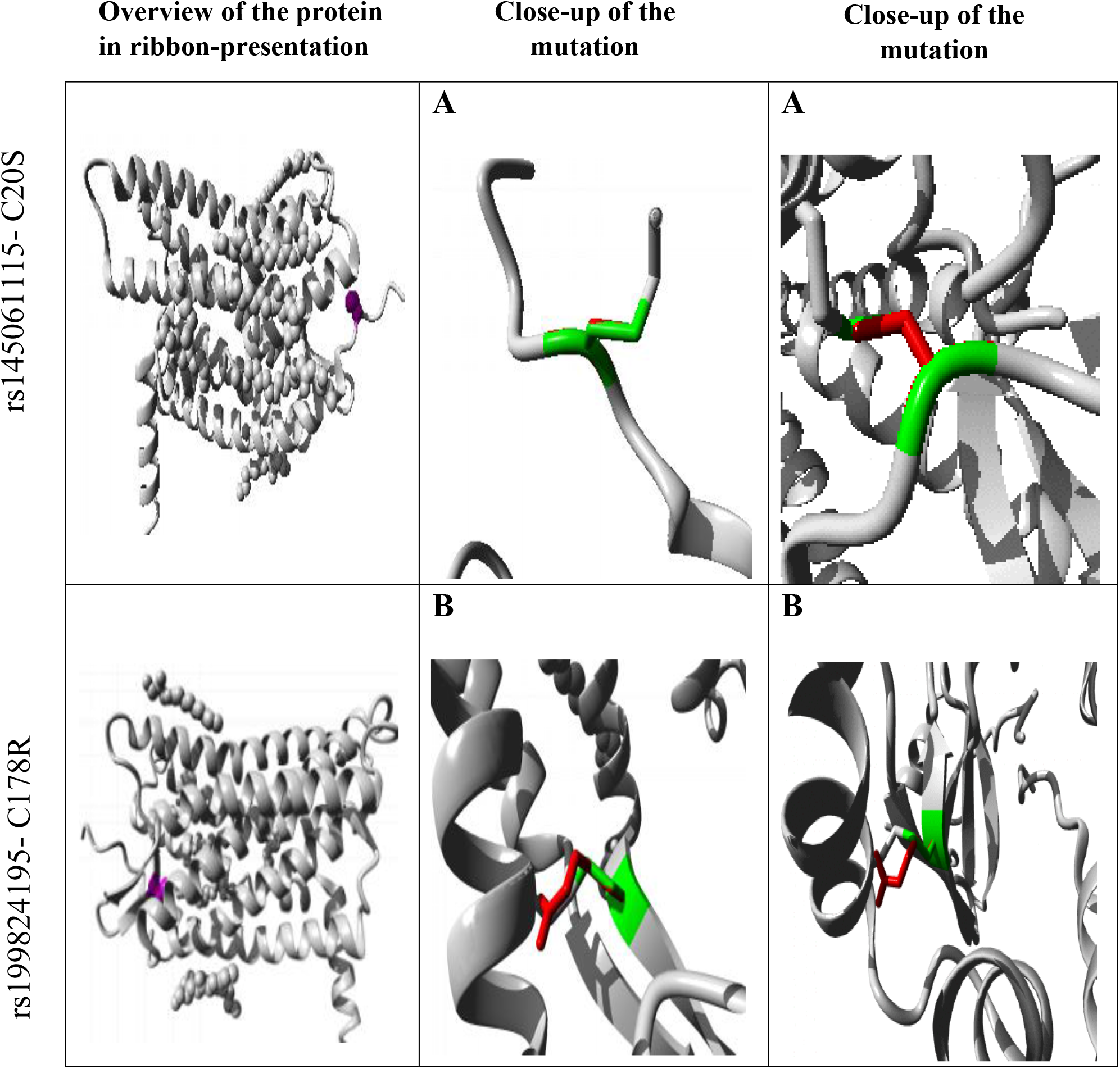
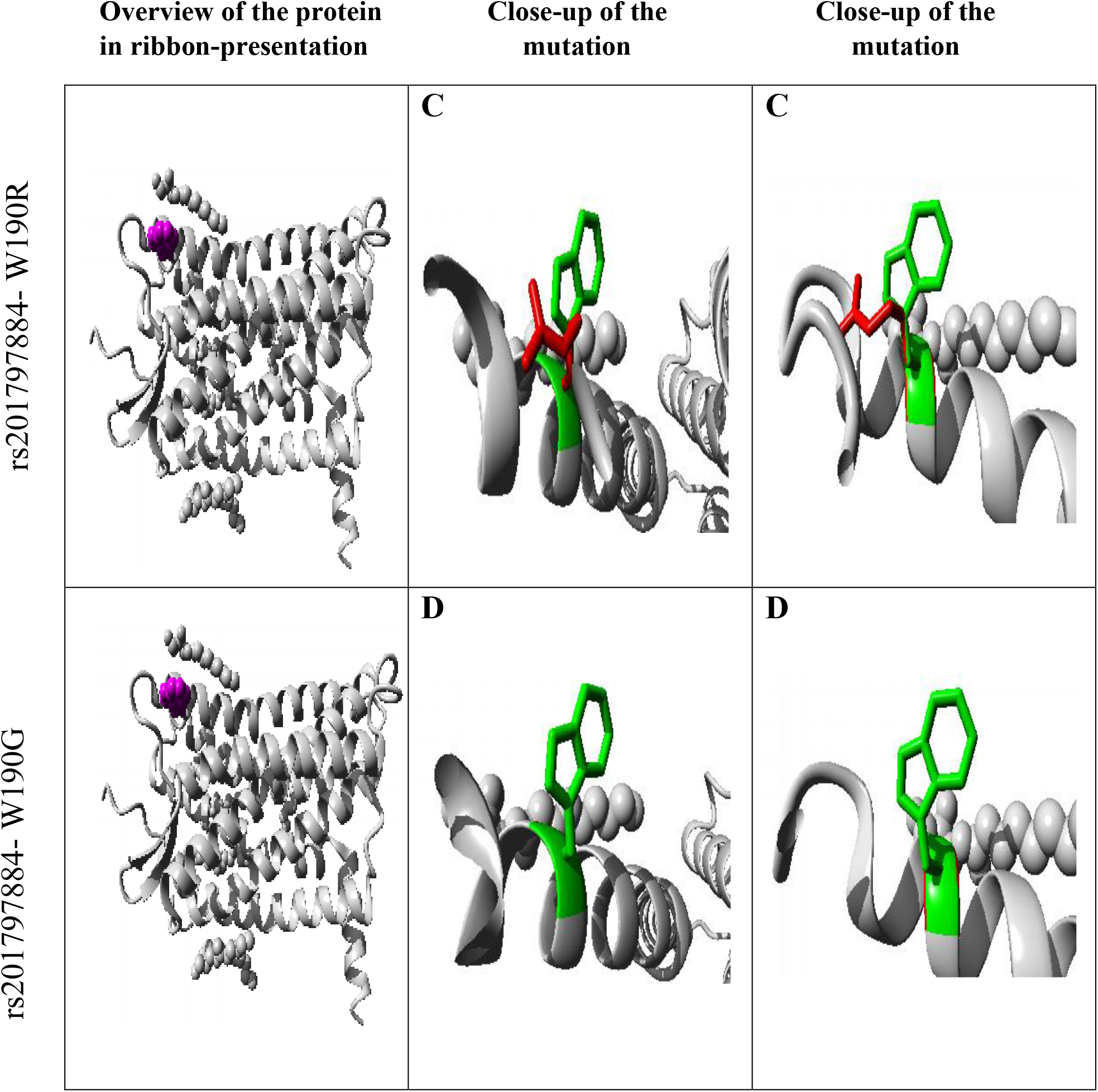
3D model of rs145061115, rs199824195 and rs201797884 show the effect of each SNP on the structure and function of protein domains. Overview of the protein in ribbon presentation shows the protein in gray and the affected amino acid as small balls in purple. In figure (2): A, B, C and D is a close-up of the mutations show the side chain of the wild type colored in the green and mutant residue colored in red for the rs145061115, rs199824195 and rs201797884 at position 20(from Cysteine to Serine),178(from Cysteine to Arginine),190(from Tryptophan to both Arginine and Glycine) respectively.

**Table 2:**
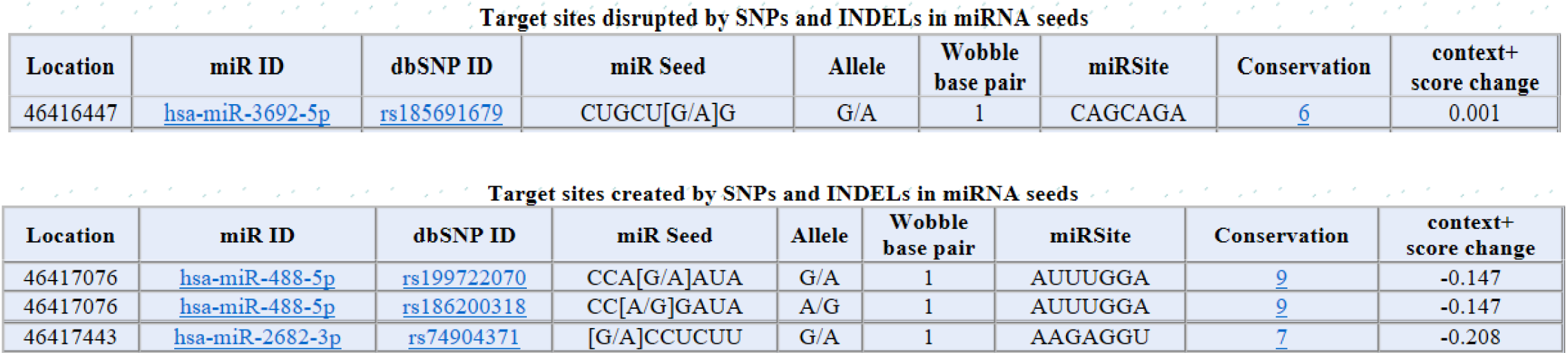
The effect of SNPs in 3’ region of CCR5 gene created by PolymiRTS show the target sites disrupted and created by SNPs in miRNA seed along with conservation score (CS) and context +score change, the higher (CS), the more profound effect is predicted to be, the higher context +score the more likelihood of change in target site of miRNA.

one miRNA [hsa-miR-3692-5p](http://microrna.sanger.ac.uk/cgi-bin/sequences/mirna_entry.pl?acc=) (with CS 6) which affected by the SNP rs185691679 are worth noting to disrupted the target site in miRNA seed.

Two miRNAs [hsa-miR-488-5p](http://microrna.sanger.ac.uk/cgi-bin/sequences/mirna_entry.pl?acc=) (with CS 9 and 7) which affected by the SNPs rs199722070 and rs186200318 respectively are worth noting to create the target site in miRNA seed.

SNPs within CCR5 gene upstream region predicted by RegulomeDB which allow focusing on noncoding variants that are likely to directly affect binding (Category 1 and 2). Category 2(a–c) demonstrates direct evidence of binding through ChIP-seq and DNase with either a matched PWM to the ChIP-seq factor or a DNase footprint this result in 3 SNP with location (chr3:46414196 (n/a), chr3:46414176 (n/a) and chr3:6414175 (n/a)) in (2a) regulome DB score, which corresponded to Likely affect binding. 39 SNPs found within in category (2b). and 6 SNPs in (2c). The rest of SNP is found to be in Category 3(a–b) which considered less confident in affecting binding and Categories 4–6 which lack evidence of the variant actually disrupting the site of binding.

**Table 3:**
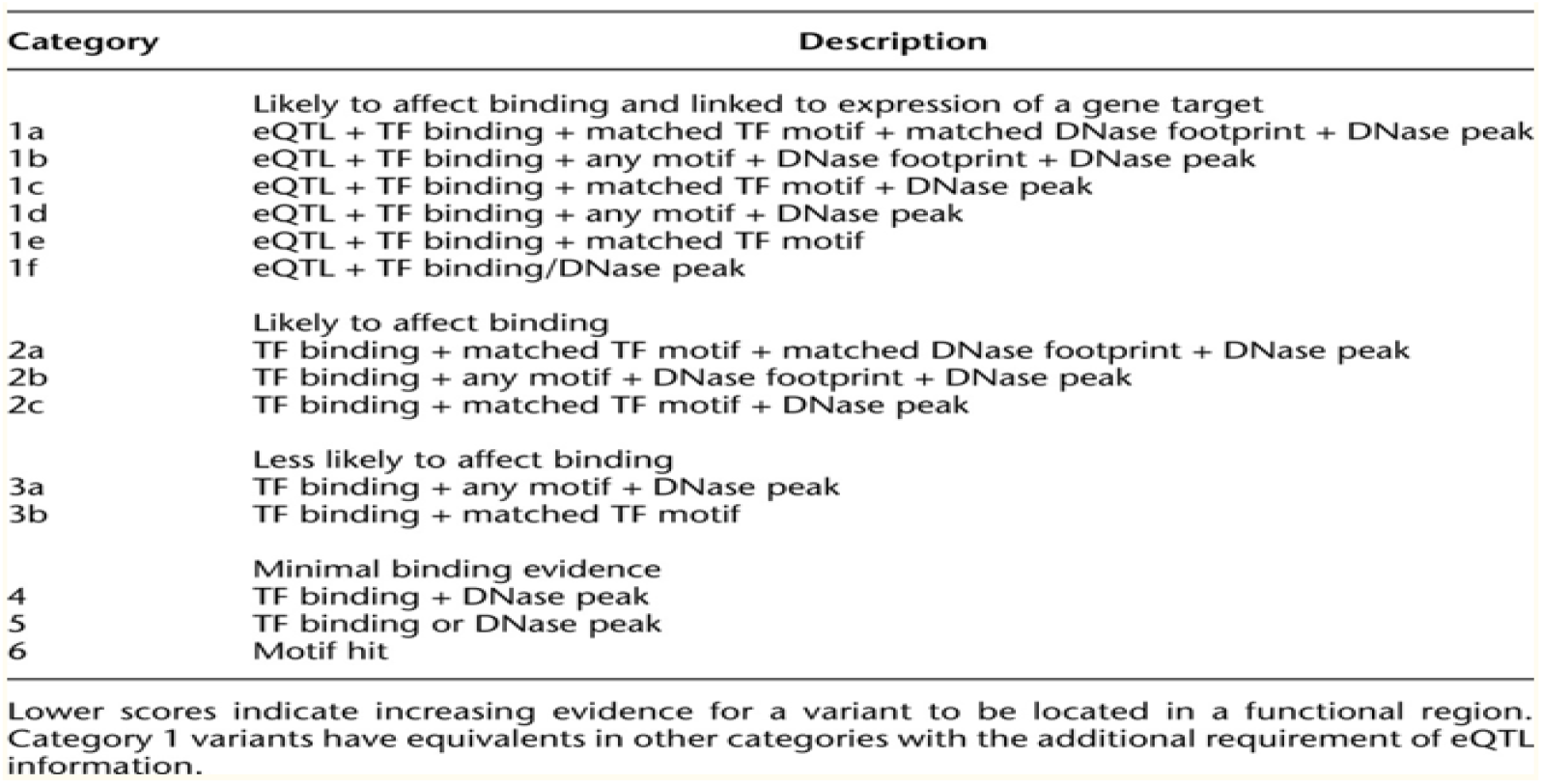
RegulomeDB variant classification scheme:

## 4. Discussion

The aim of this study was to analyze SNPs Identified in CCR5 gene using different bioinformatics tools to predict the effect of SNPs on the related protein. three (3) variant in the CCR5 binding domain with viral gp120 (rs145061115, rs199824195 and rs201797884) which correspond to C20S, C178R and (W190R/W190G) respectively lead to change in structure and function of the protein. The C20S variant located in the N-terminal in a domain that is important for the activity of the protein and in contact with residues in another domain, the interaction between these domains could be disturbed by the mutation, which might affect the function of the protein. This variant also results in destabilization of the structure because the hydrophobicity of the wild-type and mutant residue differs, this mutation might cause loss of hydrophobic interactions with other molecules on the surface of the protein, this SNP is only found in European population with minor allele frequencies (0.001%). The C178R variant, initially discovered in the Vietnamese population(23), affects a highly conserved cysteine involved in disulfide bonding between ECL-1 and ECL-2, which is important for CCR5 structure and in HIV binding(24), the structural information of this variant result from project Hope web server show that the mutant residue is bigger than the wild-type residue, the wild-type residue charge was neutral but the mutant residue charge is positive-which can lead to protein folding problems- and the wild-type residue is more hydrophobic than the mutant residue this will cause loss of hydrophobic interactions in the core of the protein. this SNP is only found in East Asian population with minor allele frequencies (0.003%). The change of a Tryptophan into an Arginine at position 190 means that the mutant residue is smaller than the wild-type residue and the wild-type residue is more hydrophobic than the mutant residue. The mutated residue is not in direct contact with a ligand, however, the mutation could affect the local stability which could affect the ligand-contacts made by one of the neighboring residues. This will cause a possible loss of external interactions.

The mutation of a Tryptophan into a Glycine at position 190 share much structural information including the size and hydrophobicity with (Tryptophan into an Arginine mutation at the same position) nevertheless this mutation introduces a glycine at this position, Glycine is very flexible and can disturb the required rigidity of the protein at this position. The minor allele frequencies of this SNP in the African and American population are 0.003 % and 0.0317% respectively. Martinson et al. analyzed the distribution of the Δ32 mutation in more than 3000 individuals from various countries and found a 2–5% gene frequency in Europe, Middle East and some parts of the Indian subcontinent (18). Isolated incidences of Δ32 found in other regions were attributed to European gene flows into these areas.

These mutations may explain and represent the molecular basis for the resistance to HIV infection that reported in many people. Because these mutations change the properties and function of the protein.

The other SNPs found in this study that affect transcription factor and miRNA binding; some are predicted to have significant effect on miRNA binding such as hsa-miR-3692-5p and hsa-miR-488-5p due to their high conservation score, and the other such as SNPs chr3:46414196, chr3:46414176 and chr3:6414175 are likely to affect transcription factor binding.

